# Application of 3D Zernike Descriptors in Antibody Structural Clustering and Repurposing

**DOI:** 10.64898/2026.08.12.744489

**Authors:** Diego da Silva de Almeida, Aline Oliveira Albuquerque, Antônio Marcelo Peixoto Lima, Eduardo Menezes Gaieta, Julia Silva Souza, Andrielly Henriques dos Santos-Costa, Luca Milério de Andrade, Jean Vieira Sampaio, Geraldo Rodrigues Sartori, e João Hermínio Martins da Silva

## Abstract

Antibodies generally exhibit high specificity for their cognate epitopes, but structural and physicochemical similarities between distinct epitopes can enable an antibody to recognize different antigens, resulting in cross-reactivity. This property can be exploited for antibody repurposing. To identify epitopes that share such similarities, both sequence- and structure-based approaches can be employed. In this context, 3D Zernike descriptors provide a compact representation of protein surface geometry as numerical feature vectors, enabling quantitative comparisons independently of structural alignment and orientation. Thus, this study aimed to evaluate the application of 3D Zernike descriptors for the structural clustering of antibodies and epitopes and to explore their use in antibody repurposing for the recognition of new targets. To this end, antibody binding sites previously associated with recognition of similar epitopes were analyzed at different structural levels, considering the CDRs, CDRH3, and complete paratopes. Surface similarity was subsequently quantified by calculating the Euclidean distance between their corresponding 3D Zernike feature vectors. Performance was benchmarked against SPACE2. Additionally, different distance thresholds were evaluated based on their ability to recover antibody pairs recognizing the same epitope. The paratope-based approach provided the best balance between the number of identified pairs and precision at a distance threshold of 2.7, whereas epitope clustering showed robust performance up to a distance of 3.0. At these thresholds, the 3D Zernike descriptors identified a greater number of functional pairs than SPACE2 while maintaining comparable precision and identifying complementary sets of antibody pairs.. BTaken together, these findings support the use of 3D Zernike descriptors for structural clustering of antibodies and epitopes and for guiding antibody repurposing G, a highly lethal zoonotic pathogen. Structural screening identified three antibodies with epitopes similar to the NiV target that also showed a consistent binding preference for the target epitope in molecular docking assays. Notably, one candidate, originally directed against a SARS-CoV-2 epitope, formed a stable complex with the NiV epitope, remaining within the 5 Å RMSD threshold during heated molecular dynamics simulations and emerging as a potential cross-reactive candidate.These results support the use of this computational framework for biopharmaceutical discovery against emerging targets. Taken together, these findings support the use of 3D Zernike descriptors for structural clustering of antibodies and epitopes and for guiding antibody repurposing.

## 1 Introduction

Antigens are molecules that bind to components of the immune system, such as B-cell (BCR) and T-cell receptors (TCR). They are also referred to as immunogens when they can additionally trigger an immune response. Antigen neutralization is mediated by antibodies, which are specialized proteins that recognize and bind with high affinity to specific regions on the antigen surface, known as epitopes. This interaction can block antigen binding to its molecular target or recruit other components of the immune system, resulting in a more efficient response (1,2). The Fab region (fragment antigen-binding) comprises both constant and variable domains, with the latter forming the Fv region that mediates antigen recognition (3). Within the Fv, binding specificity and affinity are primarily governed by the complementarity-determining regions (CDRs), which consist of three hypervariable loops from the light chain and three from the heavy chain that together shape the antigen-binding interface (4).

Despite the high specificity typically associated with antibody-epitope recognition, antibodies may exhibit cross-reactivity, defined as the ability to recognize and bind epitopes distinct from the original target antigen. A major driver of this phenomenon is epitope similarity, which may occur among proteins from the same family and across evolutionarily distinct proteins (5,6). When such cross-reactivity does not result in deleterious interactions with human self-antigens, which in many cases can trigger autoimmune responses, it plays a fundamental role in the immune system by contributing to resistance against multiple antigens, facilitating the neutralization of pathogens similar to those responsible for previous infections (7,8).

Structure and sequence-based analyses are commonly used to assess antigen similarity *in silico*. Conventional approaches include sequence alignment, aiming to identify linear segments of critical and conserved residues, or the application of atomic overlap metrics, such as RMSD (Root Mean Square Deviation) and TM-align. Strategies that rely on structural comparison are particularly relevant because similar sequences may adopt distinct conformations, whereas divergent sequences can converge toward comparable three-dimensional arrangements. Nonetheless, it is widely recognized that protein function is closely related to surface properties, including geometric, physical, and chemical features that extend beyond sequence similarity or structural overlap. As a result, protein surface representation has increasingly been incorporated in the study of geometric complementarity and protein similarity, being a cornerstone of softwares such as MaSIF, SURFACE-ID, and FP-Zernike (9–11).

However, current sequence and structure based methods present important limitations. Recent studies indicate that widely used structural frameworks, such as SPACE2, exhibit low sensitivity in identifying functional antibody pairs, largely due to constraints that include the dependence on CDR length, the availability of similar structures in the Protein Data Bank (PDB), and the reliance on atomic overlap (12). Among the current approaches, protein surface representation using feature vectors stands out by combining computational efficiency and accuracy, enabling similarity assessment through vector-based metrics. In this context, 3D Zernike descriptors provide a mathematical method capable of robustly capturing protein shape and geometric properties. They could provide an alternative strategy for capturing the structure similarity required for clustering by representing geometric surface features in an alignment-independent manner, as they are rotation-invariant and independent of sequence length (11,13).

This method has been successfully applied to several immunology-related tasks, including antibody classification according to antigen type and paratope prediction (14,15). Despite their potential, the widespread application of 3D Zernike descriptors has been limited by the computational cost and difficulties in large-scale automation. Addressing these limitations, the recently developed Python-based FP-Zernike software provides an efficient and scalable framework that facilitates protein surface representation and integrates the ability to compute similarity through Euclidean distance calculation (16).

Building on these advances, the present study aims to explore 3D Zernike descriptors in two immunoinformatics applications: the structural clustering of antibodies and epitopes, and antibody repurposing for therapeutic development. The clustering of functional antibodies that recognize the same epitope is essential for reducing the sampling space and improving analytical efficiency. The second application explores antibody repurposing, leveraging the ability of antibodies to recognize structurally related targets and enabling the repositioning of known antibodies for new clinical indications. By exploiting structural similarities between epitopes, this strategy can significantly accelerate therapeutic development through the identification of pre-existing antibody scaffolds that already exhibit a high degree of geometric and physicochemical complementarity.

Here, we propose the use of 3D Zernike descriptors to identify antibodies with potential multi-target recognition, facilitating the redirection of existing antibodies in emergency scenarios or toward clinically relevant targets. Using the Nipah virus (NiV) as a case study, we applied our repurposing workflow to propose a novel antibody targeting a key epitope of the G glycoprotein. Given that NiV is a highly lethal pathogen, often requiring quarantine measures and listed by the World Health Organization as a priority threat with pandemic potential, the prolonged timelines often required for traditional de novo discovery can hinder rapid-response efforts. This context warrants the use of 3D Zernike descriptors to rapidly redirect proven antibody scaffolds toward emergent viral targets.

## 2 Methods

### 2.1 Structural clustering of antibodies and epitopes

#### 2.1.1 Data preparation

This study utilized a dataset of functional antibodies previously curated by Waury et al. (2024). This dataset comprises 54 unique antibodies which, when paired based on their shared specificity for the same epitope on a given protein antigen, result in 256 distinct antibody pairs (17). To enable a direct comparison with SPACE2, an antibody clustering tool that requires modeled structures, structural modeling was performed (12). The sequences and experimental structures of the Fv domains in complex with their respective antigens were retrieved from the PDB. These 54 unique antibodies were then modeled using a local installation of AbodyBuilder2 integrated into the ImmuneBuilder suite, following the IMGT numbering scheme (18). From the resulting models, individual files were generated for each antibody containing: (i) all CDR residues, (ii) only the CDRH3, and (iii) the paratope. CDR residues were identified via sequence numbering using ANARCI (19) (Figure 1).

**Figure 1.**
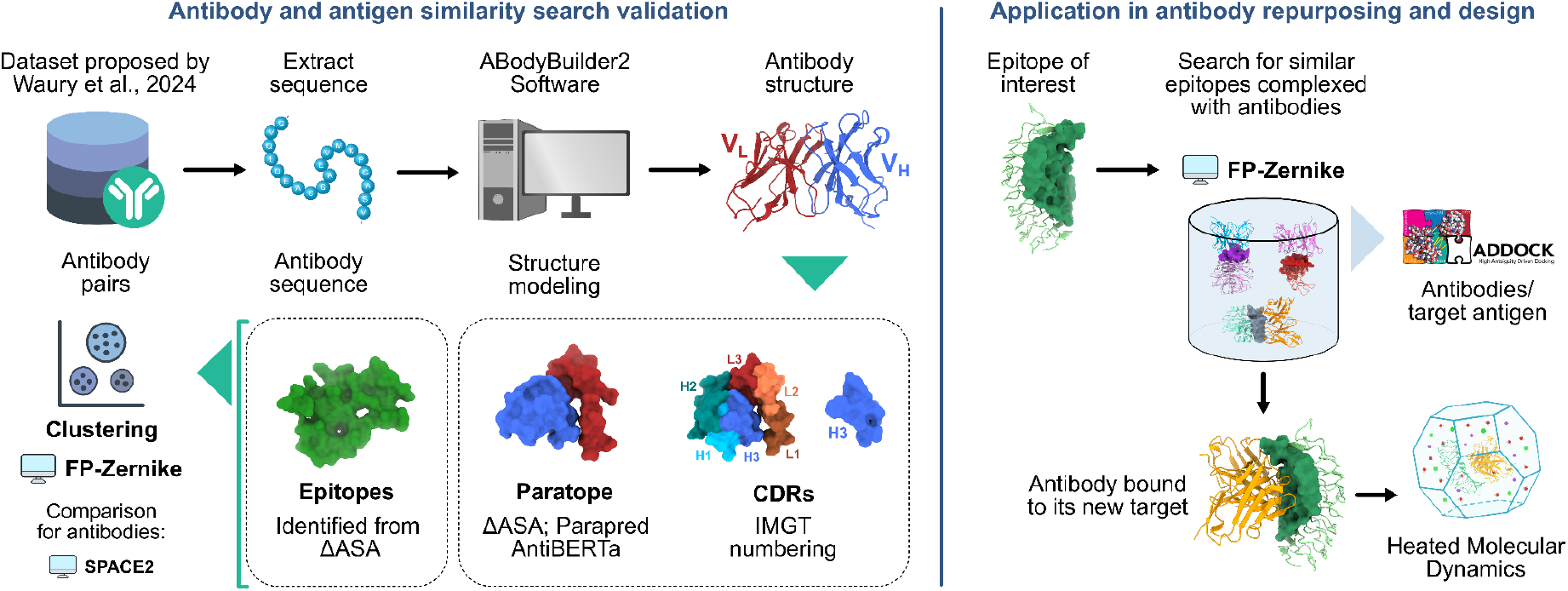
Methodological workflow highlighting the validation of antibody–antigen similarity search and its application to antibody repurposing and design. On the left, the dataset proposed by Waury *et al*. (2024) was used to extract antibody sequences, which were structurally modeled using ABodyBuilder2. The resulting structures were analyzed to identify epitopes, paratopes, and CDRs. The structural similarity among antibodies was assessed using the FP-Zernike method and compared with SPACE2. On the right, FP-Zernike is applied to identify structurally similar epitopes in antibody–antigen complexes, enabling antibody repurposing for new targets. The complexes were predicted by docking (HADDOCK 2.4), and their stability was analyzed using heated molecular dynamics simulations.

Paratope and epitope residues were defined based on changes in the solvent-accessible surface area (ΔASA) between the unbound antibody and antibody–antigen complex. Residues showing an ASA variation greater than 1 Å^2^ were considered part of the paratope or epitope (20). This approach was enabled by the availability of antibody–antigen complex structures. In scenarios where such complexes are unavailable and only antibody structures are known, paratope prediction tools, such as Parapred and AntiBERTa, have been employed (21).

For Parapred, the sequences of each CDR were extended by two flanking residues at each end. The software assigns a score to each residue, and those with scores ≥ 0.67 (default cutoff) were considered to be part of the paratope. In contrast, AntiBERTa uses the full antibody sequence and directly returns residues predicted to be part of the paratope. Based on these different strategies for representing antibodies and antigens, six PDB files were generated for each of the 54 complexes, corresponding to (i) all CDRs, (ii) CDRH3 only, (iii) paratopes defined by ΔASA, (iv) epitopes defined by ΔASA, (v) paratopes predicted by Parapred, and (vi) paratopes predicted by AntiBERTa.

#### 2.1.2 Surface representation and similarity analysis

The PS (protein surface) mode of FP-Zernike was used for structural surface representation. FP-Zernike is an open-source toolkit that efficiently computes 3D Zernike descriptors from PDB files and provides an Euclidean distance metric for quantifying surface similarity (11). According to the FP-Zernike authors, distances ≤2.5 denote similar surfaces. Given that antibody binding sites are composed of highly structurally variable loops, different cutoff values were investigated. Descriptors were calculated for all regions of interest, and similarity was evaluated for distances ranging from 2.0 to 3.0 in increments of 0.1, allowing the identification of a more appropriate cutoff for antibodies. Each identified pair was subsequently evaluated, and the recovery capacity of the methods was measured through precision. For antibody clustering using SPACE2, the modeled structures were used as inputs. Precision was calculated as the ratio of correctly identified positive pairs to the total number of pairs predicted by the method. Two clustering modes were tested, considering the full set of CDRs and only the CDRH3. In both cases, a default RMSD cutoff of 1.25 Å was applied.

### 2.2 Antibody repurposing

#### 2.2.1 Construction of the epitope set and geometric representation of its surfaces

To characterize the antigenic surfaces, 6,570 antibody-antigen complexes were retrieved from AbSet (22). This database was selected for providing a standardized set of variable region structures with uniform nomenclature. From these complexes, epitope residues were extracted using the ΔASA, constituting what we will refer to as the AbSet-Epitope Dataset. Subsequently, FP-Zernike was employed to compute three-dimensional Zernike descriptors in PS mode for each epitope. This structural characterization enabled the case study application, aimed at identifying antibodies with potential cross-reactivity through a repurposing workflow.

#### 2.2.2 Identification for structurally similar epitopes

The Nipah virus glycoprotein G (NiV-G) is essential for viral entry into host cells and represents a primary target for therapeutic antibodies. Its three-dimensional structure was retrieved from the PDB (ID: 8XPY), representing the protein in complex with a neutralizing nanobody bound to a critical and highly conserved epitope. This target epitope was defined and represented via 3D Zernike descriptors as previously described. Using the Euclidean distance metric implemented in FP-Zernike, structurally similar epitopes were searched within the AbSet-Epitope Dataset. Antibodies bound to epitopes with a Euclidean distance lower than 3.0 were selected for subsequent evaluation. These candidates were further assessed regarding their binding mode to the target epitope, as well as their complementarity and structural stability (Figure 1).

#### 2.2.3 Evaluation of the interaction mode between antibodies and target epitope

Using the antibodies identified in the previous step, molecular docking calculations were performed to obtain the binding modes of the corresponding antibody–antigen pairs. HADDOCK 2.4 was employed because of its well-established performance in antibody–antigen systems (23). Biomolecule preparation included residue renumbering to avoid overlaps, renaming of antibody and antigen chains as A and B, respectively, and defining active and passive residues. The histidine protonation states were determined using MolProbity, which is integrated into HADDOCK. For antibodies, residues belonging to the CDRs were considered active, whereas for antigens, all solvent-accessible residues, identified with FreeSASA, were defined as passive (24). In this way, a blind docking strategy was adopted aiming to evaluate whether the antibodies recognize the target epitope in an unconstrained search. The HADDOCK protocol consists of three stages: (i) randomization of initial orientations and rigid-body minimization (it0), (ii) semi-flexible refinement (it1), and (iii) refinement in explicit solvent (itW), in which 1000, 250, and 250 poses were generated, respectively. The structures obtained in the previous stage were subjected to HADDOCK clustering analysis to identify antibodies with a high binding propensity for the target epitope. Candidates were prioritized if they yielded highly populated and energetically favorable clusters, indicating a converged and stable docking solution at the interface.

Based on these results, directed docking was performed for the antibodies that demonstrated such convergence, focusing exclusively on the epitope residues and the antibody CDRs. The resulting poses were evaluated using the REF15 energy function from the Rosetta software suite, and the best-ranked configurations were subjected to structural stability evaluations.

#### 2.2.4 Stability analysis of antibody-antigen complexes

To assess complex stability, heated molecular dynamics simulations were performed using AMBER24 with up to three replicates. Each replicate was initiated from the same docking pose but with different initial random velocity assignments to ensure statistical sampling. This strategy reduces the likelihood of docking-derived false positives, as incorrect complexes tend to destabilize at elevated temperatures, whereas the correct interactions remain stable (25,26). During system preparation, the structures were protonated at pH 7.4 using PDB2PQR (27). Terminal capping was applied using acetyl (ACE) and N-methylamine (NME) groups to the antibody and antigen N-terminus and C-terminus, respectively. Subsequently, topological parameters were generated with the tLEaP application from AmberTools24 using the ff19SB force field. The system was solvated in an OPC water box with a minimum distance of 10 Å between the protein and the box edge, neutralized with Na^+^ and Cl^-^ counterions, and had its ionic strength adjusted to 150 nM. In addition, ParmEd was used for hydrogen mass repartitioning (HMR) of the topologies, allowing a 4 fs timestep during simulations.

Prior to production dynamics, energy minimization and system equilibration was performed. Initially, two stages of minimization were executed. In the first stage, only the solvent was minimized while keeping the solute restrained; this step consisted of 10,000 cycles, with the first 1,000 steps using the steepest descent algorithm followed by 9,000 steps of conjugate gradient. In the second stage, the entire system was minimized for an additional 10,000 cycles using the conjugate gradient algorithm. This was followed by a heating phase in which the system was gradually heated from 0 to 310 K under positional restraints of 100 kcal/mol·Å^2^ in the NVT ensemble using a Langevin thermostat. Subsequently, two stages of water density equilibration using a Monte Carlo barostat to maintain the system at 1 atm were performed using the NPT ensemble, starting with strong restraints of 100 kcal/mol·Å^2^ to allow adjustment of solvent volume and density, followed by weaker restraints of 10 kcal/mol·Å^2^ on the solute to promote greater relaxation. Next, the system underwent four equilibration stages characterized by a progressive reduction of positional restraints: 10 kcal/mol·Å^2^ on the protein backbone atoms, reduced to 1 kcal/mol·Å^2^, then 0.1 kcal/mol·Å^2^, and finally an unrestrained stage, resulting in full-system stabilization prior to production.After equilibration, each system was subjected to a 70 ns production simulation. During this phase, the system underwent stepwise temperature increases in four stages, the first 30 ns at 310 K, followed by 12.5 ns at 330 K, 12.5 ns at 360 K, and the final 15 ns at 390 K.

To evaluate the complex stability, the RMSD of the backbone atoms of the interface residues, defined as those within 8 Å of any alpha-carbon atom of the interacting partner, relative to the initial docking pose was calculated over the entire trajectory using CPPTRAJ. Complexes were considered stable if they maintained consistent RMSD values, ending the simulation below the 5 Å threshold (25).

## 3 Results

### 3.1 3D Zernike descriptors allow for precise and scalable antibody and epitope clustering

#### 3.1.1 Paratope and CDRH3 maximize the precision of antibody structural clustering

The 54 antibodies with well-defined similar pairs, selected to evaluate 3D Zernike descriptors as a metric for clustering structurally similar antibodies, were modeled using AbodyBuilder2. Root mean square predicted error (RMSPE) analysis, the residue level confidence metric reported by the software in which values close to zero indicate higher predictive accuracy, showed average values of 0.21 Å and 0.22 Å for framework residues (non-CDR) of the heavy and light chains, respectively. In the CDR regions, which are primarily responsible for antigen recognition, the average errors were 0.33 Å (CDRH1), 0.36 Å (CDRH2), 0.64 Å (CDRH3), 0.43 Å (CDRL1), 0.23 Å (CDRL2), and 0.44 Å (CDRL3). The RMSPE values were consistent with the intrinsically flexible and structurally diverse nature of these loops, which makes them inherently more challenging to predict with high precision.

Clustering based on CDRH3 alone began with the identification of 17 pairs, yielding a high precision of 0.94. As the Euclidean distance threshold was increased, the number of pairs progressively grew, reaching 28 pairs with a precision of 0.75 at a distance of 2.5 and ending with 56 pairs with a precision of 0.76 (Figure 1A). Moreover, increasing the threshold up to 2.7 led to a concomitant rise in the Jaccard index, reflecting higher epitope similarity among the identified pairs and suggesting reduced false positive inclusion.

Clustering based on the ASA-defined paratope (paratopeASA) initially detected only three clusters. However, at a Euclidean distance threshold of 2.7, 74 pairs were identified with a precision of 0.80. In cases where the paratope was predicted by Parapred and AntiBERTa, the same distance threshold of 2.7 Å provided the best balance between the number of detected pairs and precision, resulting in 43 pairs (precision of 0.81) and 45 pairs (precision of 0.80), respectively (Figure 2A). As observed for the CDRH3-based clustering approach, a distance threshold of 2.7 preserved the Jaccard index above 0.6, approaching the upper limit of 0.75 and supporting strong epitope similarity. Thus, the paratope-based approach demonstrated a balance between the number of detected pairs and precision, allowing the extension of the distance threshold without significantly compromising reliability.

**Figure 2.**
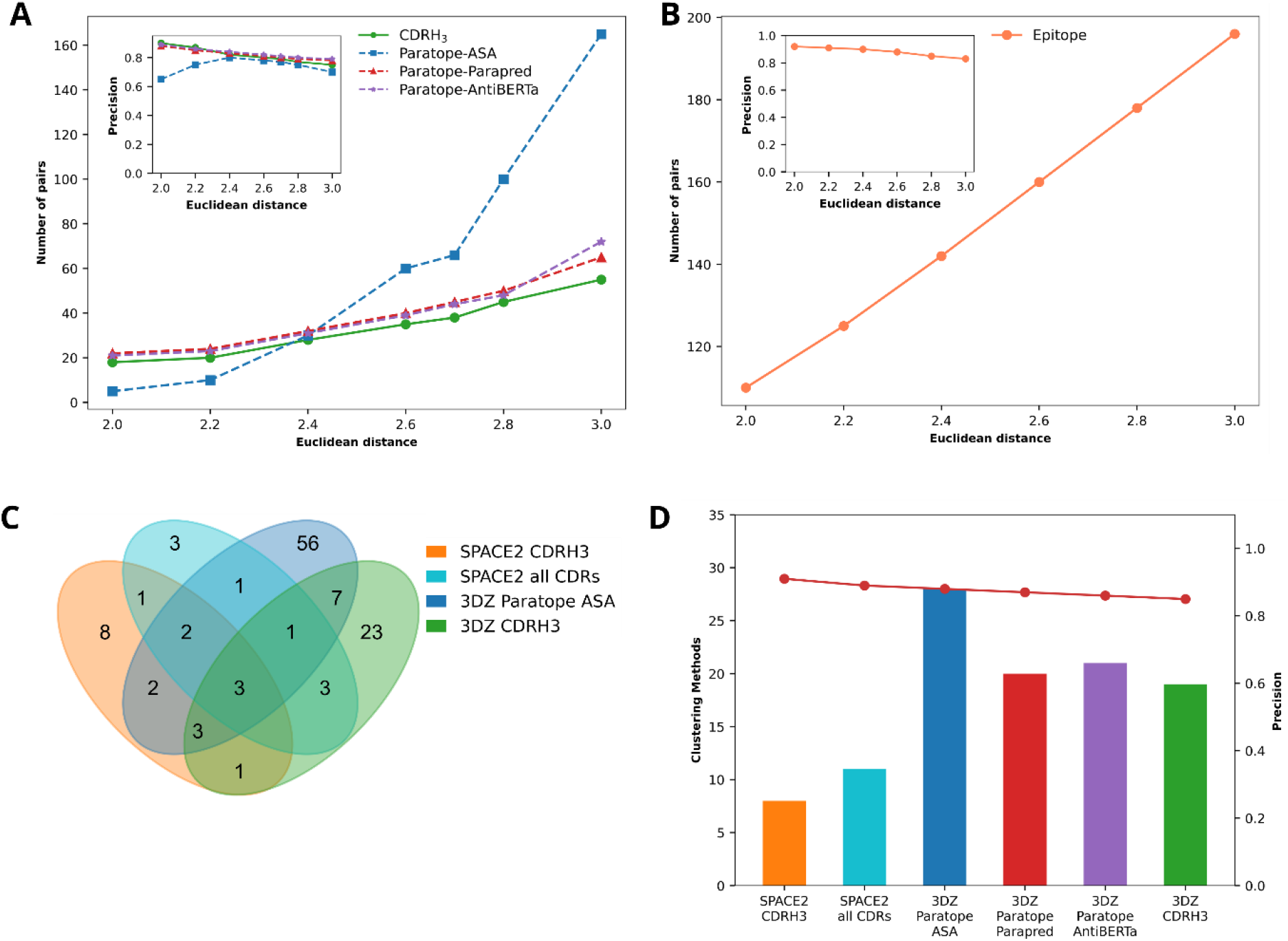
Evaluation of Antibody Clustering Approaches. (A) Comparison of different antibody clustering approaches using 3D Zernike descriptors, highlighting the number of identified pairs and their accuracy (inset). (B) Number of similar epitope pairs and accuracy (inset) obtained by clustering with 3D Zernike descriptors. (C) Venn diagram illustrating the overlap of pairs obtained by clustering with SPACE2 (considering all CDRs and only CDRH3) and with 3D Zernike descriptors, using paratopeASA and CDRH3 and a 2.7 Euclidean distance cutoff. (D) Comparison of the percentage of identified pairs (bars) and accuracy (line) between antibody pairs detected by SPACE2 (considering all CDRs and only CDRH3) and pairs identified by 3D Zernike descriptors, using different approaches: paratopeASA, Parapred and AntiBERTa.

In contrast, a distinct behavior was observed when all the CDRs were considered. A large number of pairs were identified at the outset, with 352 pairs detected immediately, followed by a steep rise to 597 pairs at a Euclidean distance threshold of 3.0. However, evaluation of the precision indicated values of 0.57 at the initial threshold, decreasing to 0.40 as the cutoff increased. This decline in precision indicates that although the all-CDR approach captures a larger number of pairs, it includes a substantial number of false positives, thereby compromising its reliability.

#### 3.1.2 Expanding the detection of functional antibody pairs

Antibody clustering performed by SPACE2 identified 18 and 22 pairs using all CDRs and CDRH3-based approaches, with precisions of 0.89 and 0.91, respectively. Compared to clustering with 3D Zernike descriptors at the established threshold of 2.7, the paratope- and CDRH3-based Zernike approaches outperformed SPACE2 in pair identification, with the paratopeASA clustering method recognizing three times more pairs while maintaining similar precision (Figure 2B). To explore the complementarity between the methods, we analyzed the overlap of pairs identified via SPACE2 and 3D Zernike descriptors. We observed that 3D Zernike descriptors applied to the ASA-defined paratope and CDRH3 detected 56 and 23 pairs, respectively, which were not identified by SPACE2, whereas SPACE2, using all CDRs and only CDRH3, identified eight and three unique pairs, respectively (Figure 2C). These results indicate that the methods exhibit complementary characteristics and can be combined to increase the number of detected interactions.

#### 3.1.3 3D Zernike descriptors identify structurally similar epitopes

To extend the analysis from antibody clustering to epitope-level similarity, we evaluated the effectiveness of 3D Zernike descriptors in identifying structurally similar epitope pairs applying FP-Zernike calculations to a AbSet-Epitope Dataset. Precision remained above 0.90 up to a distance cutoff of 2.4, corresponding to the identification of 142 antibody pairs (55.47% of the total dataset). As the cutoff increased, the precision gradually declined, reaching 0.83 at a cutoff of 3.0, at which point 196 pairs (76.56% of the total dataset) were recognized (Figure 2D). Collectively, these results indicate that the method enables robust epitope comparison and supports the use of a Euclidean distance cutoff of up to 3.0, providing an effective balance between the number of identified pairs and precision.

### 3.2 Screening based on structural similarity identifies antibodies with the potential to recognize the Nipah virus epitope

#### 3.2.1 Identification of new antibodies complementary to the Nipah Virus epitope

Building upon the robust performance of 3D Zernike descriptors in identifying structural similarities, we sought to apply this methodology to a relevant therapeutic target. We investigated, within the AbSet dataset, antibodies capable of binding to epitopes structurally similar to the epitope of the Nipah virus glycoprotein G (NiV-G). This transition from methodological validation to functional application allowed us to evaluate the practical utility of our workflow in a drug repurposing context. Screening for similar epitopes using the Zernike descriptor method resulted in the identification of four candidate structures: one human T-cell immunoreceptor, two antibodies targeting the dengue virus envelope protein E, and one antibody against the SARS-CoV-2 Spike protein. Focusing on therapeutic design, subsequent analyses were restricted to the three antibodies that did not recognize the human antigen, thereby mitigating potential safety concerns, as cross-reactivity with human targets often correlates with an increased risk of off-target effects and immunogenicity.

#### 3.2.2 Mode of interaction and stability of complementary antibodies

To investigate whether the selected antibodies exhibited a tendency to recognize the NiV glycoprotein G epitope, we analyzed the centers of the most populated clusters obtained from the blind docking with each of the antibodies. Antibodies from PDB entries 4UTB_1 and 5N09_2 had 50 and 55 structures in their most populated clusters, respectively (Figures 3A and 3B). The antibody corresponding to PDB 7YVM_1 presented 39 structures in the most populated cluster (Figure 3C). Notably, the cluster centers of all antibodies were located on the identified glycoprotein G epitope, indicating a consistent preference for this region (Figure 3D). Based on these findings, all three antibodies advanced to the subsequent stages of directed docking and heated molecular dynamics simulations.

**Table 1.**
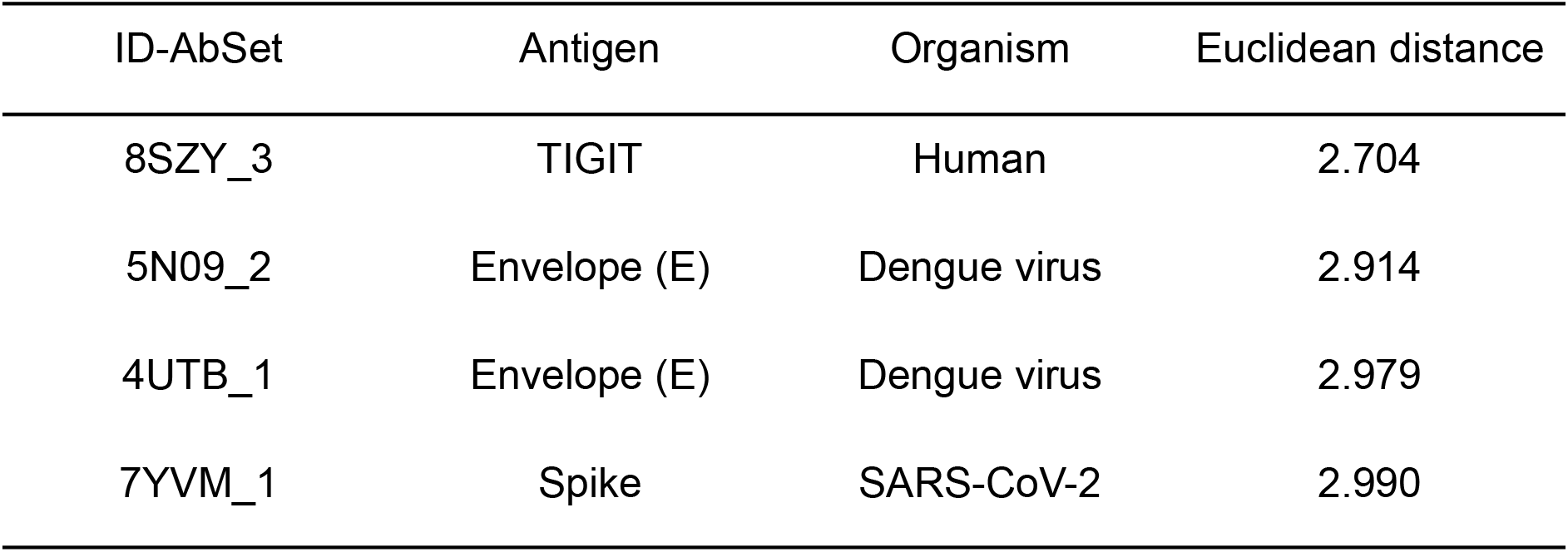
Epitopes structurally similar to the Nipah virus G glycoprotein target identified in the AbSet repertoire.

**Figure 3.**
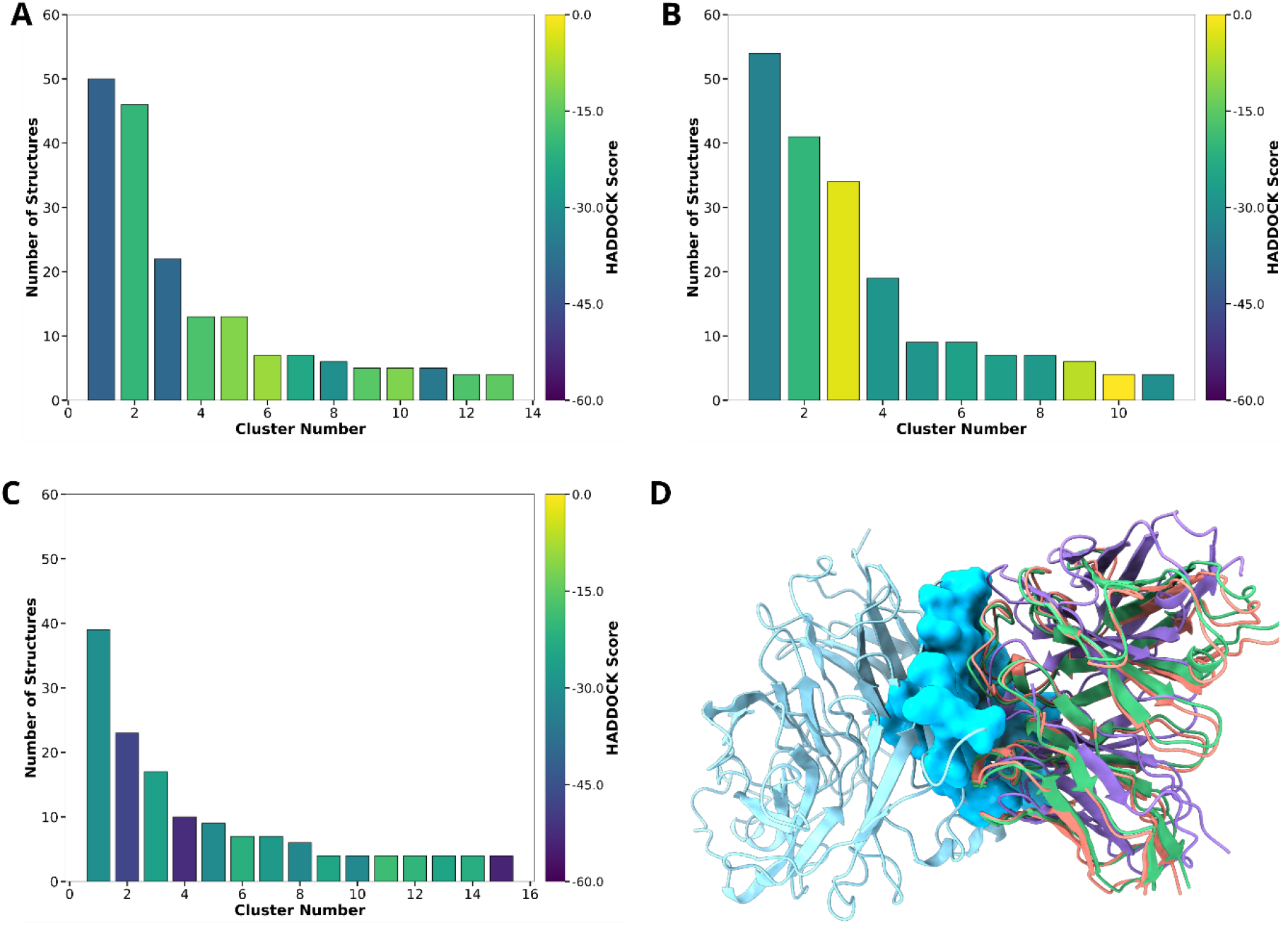
Distribution of the most populated clusters and binding mode of the cluster centers for the selected antibodies in blind docking against the Nipah virus glycoprotein G. (A) Clustering of poses for antibody 4UTB_1. (B) Clustering of poses for antibody 5N09_2. (C) Clustering of poses for the antibody 7YVM_1. The bars are colored according to the average HADDOCK score of each cluster. (D) Superposition of the centers of the most populated clusters for the three antibodies, showing that all are located on the epitope, indicating a consistent preference for the recognition of this region.

To refine the binding poses obtained from the directed docking, we performed a structural ranking using the REF15 scoring function. The top-scoring poses for each antibody yielded values of −26.10, −16.35, and −21.69 Rosetta Energy Units (R.E.U.) for PDBs 4UTB_1, 5N09_2, and 7YVM_1, respectively. While these scores provided an initial estimate of binding affinity, we subsequently evaluated the complex stability throughout heated molecular dynamics to discriminate potential false positives from truly stable interactions. Following the established protocol of performing up to three replicates only if the previous ones remained within the stability threshold, the first two antibodies—both originally directed to dengue virus antigens (4UTB_1 and 5N09_2)—did not remain stable. In the first replicate, both complexes exceeded the 5 Å RMSD cutoff at 360 K and displayed multiple instability events (Figure 4A). Consequently, these candidates were eliminated from further analysis without the need for additional replicates (Figure 4C). In contrast, the antibody corresponding to PDB 7YVM_1 remained stable across all three replicates (Figure 4B). This candidate demonstrated robust structural integrity throughout the stepwise temperature increases, without exhibiting any instability events or violating the established RMSD cutoff (Figure 4C).

**Figure 4.**
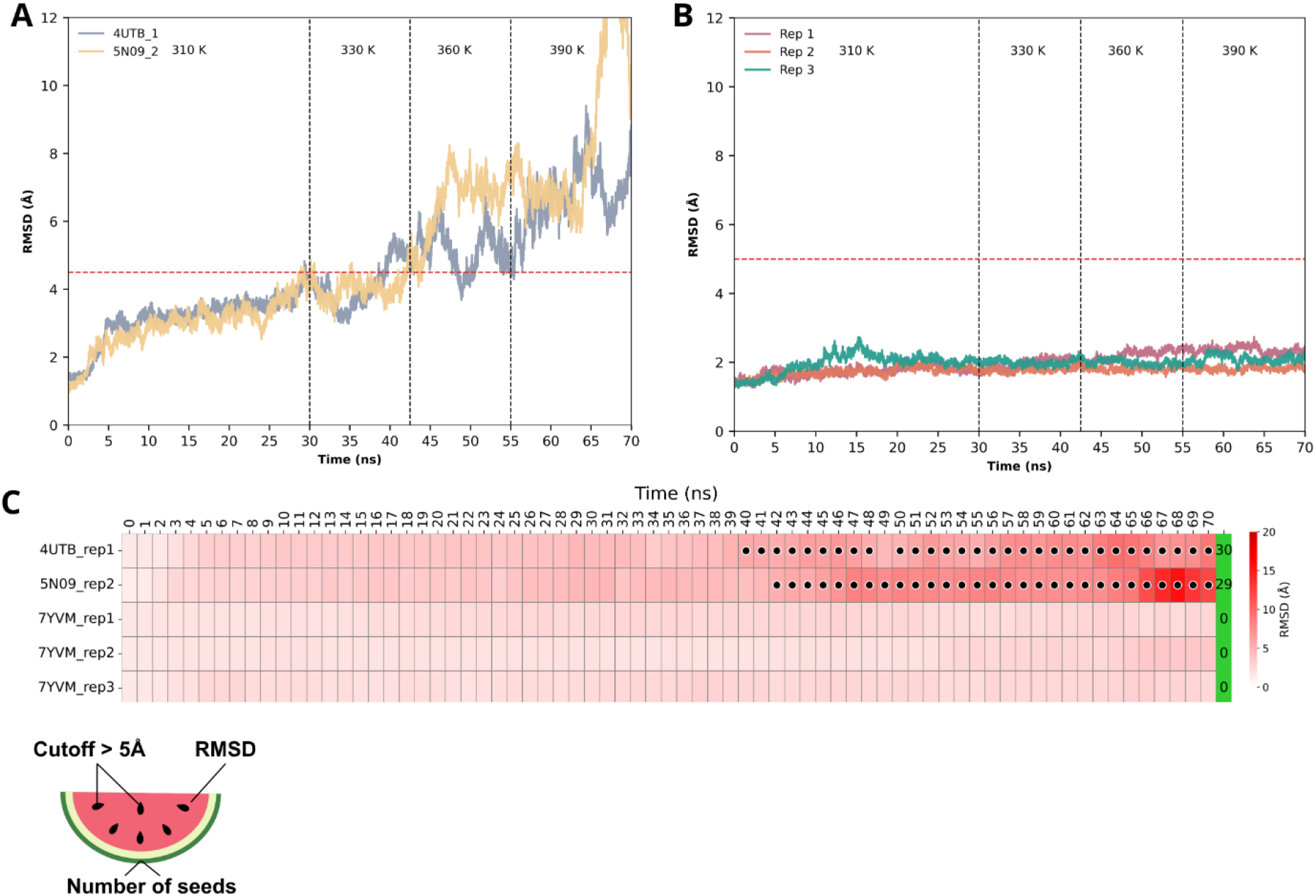
RMSD profiles of antibody–antigen complexes during heated molecular dynamics simulations. (A) Antibodies 4UTB_1 and 5N09_2 exceeded the 5 Å stability cutoff at 360 K, showing multiple instability events. (B) Antibody 7YVM_1 remained stable across all three replicates, maintaining RMSD values below the cutoff throughout the temperature increase. (C) Watermelon plot summarizing RMSD per nanosecond, where seeds indicate time points exceeding the 5 Å cutoff; a seedless profile reflects continuous complex stability.

## 4 Discussion

The higher number of retrieved similar antibody pairs using 3D Zernike descriptors may be attributed to its broader applicability, as the method is not restricted by requirements such as consistent CDR lengths imposed by SPACE2. Additionally, this method is free from atomic overlap, making the calculation faster, and introducing an approach that aligns with new protein similarity methods based on surface properties (Riahi *et al*., 2023; Ye *et al*., 2024). Among the different approaches analyzed, the clustering method based on paratope stood out as the most promising. The paratope is one of the most biologically relevant regions of the antibody, as it is responsible for the interaction with the antigen, presenting a surface that is complementary to its target. This characteristic provides a significant advantage over the exclusive use of CDR residues in antibody clustering, as demonstrated by Madsen et al., 2024, who investigated the structural trends in antibody-antigen binding interfaces. Although most paratope residues are concentrated in the CDRs, a considerable portion is located in the framework regions. Additionally, not all CDRs establish direct contact with the antigens.

The 3D Zernike descriptors also proved to be efficient in clustering protein epitopes. The search for similarity between epitopes is of utmost importance, as it can reveal insights related to the immune system, such as cross-reactivity, a phenomenon associated with structural similarity between epitopes (Lv *et al*., 2025; Buraphaka *et al*., 2024; Pretti *et al*., 2025; Uhuami *et al*., 2024). Thus, this new epitope-clustering approach with high precision can significantly contribute to research in this area.

Building upon this high-precision clustering, we applied the concept of epitope similarity searching to establish an antibody repurposing strategy, identifying a novel antibody with the potential to recognize the Nipah virus G glycoprotein. As discussed by Islam (2024), this approach holds significant potential for both preempting drug development for emerging diseases or general clinical contexts and accelerating regulatory approval processes, an aspect especially relevant to our study, given that the Nipah virus is classified by the WHO as a priority pathogen with pandemic potential. Although this strategy is promising and has already been applied in other contexts, frequently guided by cross-reactivity, as presented by Rodriguez-Quijada (2020), to the best of our knowledge, a clear computational guideline directing antibody identification for this type of repurposing had not yet been established in the literature.

## 5 Conclusion

In this study, we proposed the application of 3D Zernike descriptors as a method for clustering antibodies that bind to the same epitope. To achieve this, we suggest that the most effective approach is to use the paratope (whether extracted by ASA or predicted) with a similarity cutoff measured by a Euclidean distance of 2.7. Additionally, we demonstrated that this method is also effective for epitope clustering, with a distance cutoff of up to 3.0. Applying this validated methodology, we identified a candidate antiviral antibody with potential capacity to recognize the NiV glycoprotein G and maintain stable interactions with the target epitope, representing a promising scaffold for biopharmaceutical development.

